# A role for BCL6 in maintaining CX3CR1^+^ CD4^+^ T cells during helminth infection

**DOI:** 10.1101/2023.06.27.546755

**Authors:** Denis G. Loredan, Joseph C. Devlin, Kamal M. Khanna, P’ng Loke

**Author notes:** Correspondence: P’ng Loke; Kamal M. Khanna. Equally contributing corresponding authors (Kamal M. Khanna, P’ng Loke). Regeneron Pharmaceuticals, Inc., Tarrytown, NY 10591.

## Abstract

Distinct subsets of T lymphocytes express CX3CR1 under inflammatory conditions, but little is known about CX3CR1^+^ CD4^+^ T cells during Type 2 inflammation in helminth infections. Here, we used a fate-mapping mouse model to characterize CX3CR1^+^ CD4^+^ T cells during both acute *Nippostrongylus brasiliensis* and chronic *Schistosoma mansoni* helminth infections, revealing CX3CR1^+^ CD4^+^ T cells to be an activated tissue homing subset with varying capacity for cytokine production. Tracking these cells over time revealed that maintenance of CX3CR1 itself along with a T_H_2 phenotype conferred a survival advantage in the inflamed tissue. Single-cell RNA-sequencing analysis of fate-mapped CX3CR1^+^ CD4^+^ T cells from both the peripheral tissue and the spleen revealed a considerable level of diversity and identified a distinct population of BCL6^+^ TCF-1^+^ PD1^+^ CD4^+^ T cells in the spleen during helminth infections. Conditional deletion of BCL6 in CX3CR1^+^ cells result in fewer CX3CR1^+^ CD4^+^ during infection, indicating a role in sustaining CD4^+^ T cell responses to helminth infections. Overall, our studies revealed the behavior and heterogeneity of CX3CR1^+^ CD4^+^ T cells during Type 2 inflammation in helminth infections and identified BCL6 to be important in their maintenance.

## Introduction

Parasitic helminths are a major global health threat, infecting over 1 billion people globally and resulting in the loss of millions of disability-adjusted life years (DALYS) (1). The most common human helminth infections are caused by soil-transmitted helminths such as hookworms as well as various species within the genus *Schistosoma*, most prominently *Schistosoma mansoni* and *Schistosoma haematobium* (1). Hookworm infections commonly occur through the skin, often from walking barefoot on soil contaminated by fecal matter (2).

Schistosome infections, comparatively, also infect through the skin, but occur in contaminated water, such as a population adjacent lake or river (3). Attempts have been made to prevent helminth infections through better sanitation and education about hygiene, although these interventions are limited due to lack of economic resources. Widespread drug treatment, referred to as mass deworming, with drugs such as albendazole for hookworm infection and praziquantel for schistosome infection have been implemented. However, these treatments are not considered an adequate solution, as many people can be rapidly reinfected, and there are concerns about drug resistance (4–6). New forms of prevention and treatment are needed, in the form of effective vaccines and host-directed therapies. It is therefore crucial to develop a more thorough understanding of how the immune system responds to helminth infections at a cellular and molecular level.

CX3CL1, also known as fractalkine or neurotactin, is a unique chemokine in that it is the only member of the CX3C chemokine family discovered to date, with 3 amino acids separating the initial cysteine residues (49). Upon inflammation, endothelial and epithelial cells will express CX3CL1 as a membrane-bound protein, which forms tight interactions with immune cells expressing the cognate receptor, CX3CR1, and can mediate immune cell extravasation into the tissue (7). The metalloproteinases ADAM10 and ADAM17 can cleave the mucin-like domain, releasing a soluble form of the chemokine that serves to attract immune cells expressing CX3CR1 (8–9). Many tissue-specific roles of the CX3CL1-CX3CR1 axis have been characterized, revealing CX3CL1 to be a rather broadly expressed molecule, with different cell types expressing it depending on the tissue and inflammatory state (10). There are many different types of CD45^+^ leukocytes that express CX3CR1, including monocytes, macrophages, dendritic cells, Natural Killer (NK) cells, and T and B lymphocytes (11–18). The binding of CX3CL1 to its receptor has been associated with inducing behavior changes in the cell, such as altered trafficking, but also in downstream signaling resulting in increased cellular survival (19–22).

Studies on CX3CR1 biology has focused on its expression on monocytes, however both CD4^+^ and CD8^+^ T lymphocytes have the capacity to express CX3CR1 upon stimulation and activation (17). Recent work on CD8^+^ T cells showed that CX3CR1 expression could differentiate different subsets of memory T cells during viral infection, with one study finding that in mice, following infection with lymphocytic choriomeningitis virus (LCMV) and subsequent clearance, CX3CR1 expression divided memory CD8^+^ T cells into 3 groups: CX3CR1^+^ effector memory (T_EM_), CX3CR1^-^ central memory (T_CM_), and a newly proposed subset of CX3CR1^Int^ cells termed peripheral memory (T_PM_) that surveilled the tissue (23).

Another study on murine CD8^+^ T cells showed similar results, with T_EM_ cells being defined as CX3CR1^+^ following infection with *Listeria monocytogenes* (24). However, there might exist significant variation within the T_EM_ population, as one study indicated that this population included a further subset termed terminal-T_EM_ to indicate a more terminally differentiated state, that uniquely expressed CX3CR1 (25). CX3CR1 on CD4^+^ T cells has been studied less extensively than that of CD8^+^ T cells, with some studies showing CX3CR1^+^ CD4^+^ T cells to exhibit a similar phenotype to CX3CR1^+^ CD8^+^ T cells in that they appear terminally differentiated and cytotoxic (23,50). During murine LCMV infection CX3CR1 expression on CD4^+^ T cells was contingent upon expression of the costimulatory receptor GITR, and marked a population of terminally differentiated, antigen-specific T_H_1 cells that express high levels of granzyme and perforin relative to other CD4^+^ T cells, consistent with a cytotoxic phenotype (26). During *Mycobacterium tuberculosis* (Mtb) infection, T_H_1 cells could be divided into two subsets based on expression of CX3CR1, or CXCR3 (27). Interestingly, the CX3CR1 subset was preferentially located in the lung vasculature rather than in the parenchyma, and contributed little to infection control (27). Along with the LCMV study which found that CX3CR1 itself did not contribute to CD4^+^ T cell accumulation in the tissue, these data suggest CX3CR1 expression does not always result in endothelial translocation, but instead might serve as marker of the most terminally differentiated CD4^+^ T cells, lacking the plasticity need to respond to a dynamic infection. It is unclear if CX3CR1^+^ T_H_1 cells are a transitory cell type between effector and memory, or whether they represent the final stage of a particular trajectory, that may or may not aide in immunity depending on the context.

While the Type 1 inflammatory response to intracellular bacteria and viruses is characterized by the development of T_H_1 cells, the Type 2 inflammatory response to helminth infections induces CD4^+^ T cell differentiation to a T_H_2 phenotype. One study found that CX3CR1^+^ T_H_2 cells are pathogenic in a mouse model of allergic asthma, where they contribute to pathology by increasing Type 2 inflammation (28). While CX3CR1 itself was dispensable for T_H_2 migration to the lung, it did convey a survival advantage to the cells that expressed it, resulting in less apoptosis. This phenotype could be rescued in CX3CR1 deficient CD4^+^ T cells that were transduced with the survival factor BCL-2, consistent with the notion that CX3CR1 signaling is anti-apoptotic. CX3CR1^+^ T_H_2 cells were further characterized in a mouse model of atopic dermatitis, where they could drive accumulation of eosinophils in the skin (29). Using a CX3CR1 deficient mouse, it was found that CX3CR1 was required for T cell retention in the skin, with both T_H_1 and T_H_2 cells contributing to pathology. Importantly, in both studies, CX3CR1 deficiency reduced pathology, suggesting the CX3CL1-CX3CR1 axis could be targeted therapeutically in allergic disease (28,29). However, these results also indicated that CX3CR1 could be playing different roles depending on the disease and tissue affected, such as contributing to T cell survival or involvement in tissue transit.

In this study, we utilized a tamoxifen inducible fate-mapping mouse model to track CX3CR1^+^ CD4^+^ T cells during either acute *Nippostrongylus brasiliensis* or chronic *Schistosoma mansoni* infection. We find that CX3CR1^+^ CD4^+^ T cells accumulate primarily in the tissues affected by the helminths instead of lymph nodes and CX3CR1 may confer a survival advantage in the inflamed tissue. Single-cell RNA-sequencing (scRNA-seq) identified a distinct BCL6^+^ TCF-1^+^ PD1^+^ population and conditional deletion of BCL6 in CX3CR1^+^ cells result in fewer CX3CR1^+^ CD4^+^ cells during infection. Overall, our studies describe the behavior of CX3CR1^+^ CD4^+^ T cells in helminth infections and a role for BCL6 in their maintenance.

## Materials and Methods

### Mice

B6.*Cx3cr1*^CreERT2-IRES-EYFP/+^ were generously provided by D. Littman (Department of Pathology, Department of Microbiology, NYU Langone Health, New York, NY 10016). B6.*Rosa26*^stop-tdTomato^ mice (JAX: 007914) were from Jackson Laboratories (Bar Harbor, ME).

B6.*Cx3cr1*^CreERT2-IRES-EYFP/+^ mice and B6.*Rosa26*^stop-tdTomato^ mice were crossed to generate *Cx3cr1*^CreERT2-IRES-EYFP/+^ *Rosa26*^tdTomato/+^ mice as previously described (12). B6.129P2(Cg)- *Cx3cr1^tm1Litt^*/J mice were from Jackson Laboratories (strain #005582) (common name: CX3CR1^eGFP^) (10). C.129-*Il4^tm1Lky^*/J mice were from Jackson laboratories (strain #004190) (common name: IL-4/GFP-enhanced transcript [4get]) and backcrossed to C57BL/6J (strain #000664) (35). B6.129S(FVB)-*Bcl6^tm1.1Dent^*/J were from Jackson Laboratories (strain #023727) (common name: Bcl6^Fl^) and crossed to B6.*Cx3cr1*^CreERT2-IRES-EYFP/+^ mice to generate CX3CR1^CreERT2^ Bcl6^Flox^ mice (47). To induce labeling, Tamoxifen (Sigma-Aldrich) was dissolved in corn oil and given by oral gavage at a dose of 500 mg/kg body weight.

Alternatively, mice were placed on Tamoxifen diet (Teklad Global, 250, 2016, Red) with diet code TD130856, to induce conditional deletion. A mix of male and female mice were used for experiments. All mice were age matched within experiments and used at 8-15 weeks of age. All animal procedures were approved by the NYU School of Medicine IACUC Committee.

### Models of Helminth Infection

For experiments with *Schistosoma mansoni* infection, mice were infected percutaneously with 75 cercariae and analyzed at 8 weeks or 12 weeks post-infection. The reagent was provided by the NIAID Schistosomiasis Resource Center of the Biomedical Research Institute (Rockville, MD) through NIH-NIAID Contract HHSN272201700014I. NIH: *Biomphalaria glabrata* (NMRI) exposed to *Schistosoma mansoni* (NMRI). For experiments with *Nippostrongylus brasiliensis*, mice were infected with 625 L3 stage larvae by subcutaneous injection and analyzed at 10 days or 35 days post-inection. Mouse-adapted *N. brasiliensis* was maintained in the laboratory by serial passage intermittingly through C57BL/6J mice and STAT6-Knockout mice (31).

### Immune Cell Isolation

Liver tissues were processed as described. Livers were chopped and incubated in RPMI + 10% FBS with collagenase VIII (100U/mL; Sigma Aldrich) and DNase I (150 μg/mL; Sigma Aldrich) for 45 minutes at 37° C and then passed through a 70-μm cell strainer (Fisher Scientific).

Leukocytes were enriched for by density-gradient centrifugation over a 40/80% Percoll (GE Healthcare) gradient, and remaining red blood cells were lysed with ACK lysis buffer (Lonza) and washed in PBS and used for cell sorting or flow cytometry analysis.

Lung tissues were processed as described in a protocol for immune cell isolation from mice infected with *Nippostrongylus brasiliensis* (48). Lungs were chopped and incubated in IMDM + 10% FBS with collagenase I (2.4mg/mL; Sigma Aldrich) and DNase I (100 μg/mL; Sigma Aldrich) for 45 minutes at 37° C and then passed through a 70-μm cell strainer (Fisher Scientific). Red blood cells were lysed with ACK lysis buffer (Lonza) and washed in PBS and used for flow cytometry analysis.

Spleen tissue was processed as follows. Spleens were removed and placed in RPMI + 10%FBS. Spleens were then passed through a 70-μm cell strainer (Fisher Scientific). Red blood cells were lysed with ACK lysis buffer (Lonza) and washed in PBS and used for cell sorting or flow cytometry analysis.

Mediastinal and mesenteric lymph node tissue was processed as follows. Lymph nodes were removed and placed in RPMI + 10%FBS. Lymph nodes were then passed through a 70-μm cell strainer (Fisher Scientific) and washed in PBS and used for cell sorting or flow cytometry analysis.

### Flow Cytometry and Cell Sorting

Single-cell suspensions were stained with fluorescently conjugated antibodies in a 1:100 dilution unless otherwise noted. For live/dead discrimination, cells were stained with either LIVE/DEAD Blue Reactive Dye (Invitrogen) (20 minutes, on ice, before primary) or Alexa Fluor NHS Ester (ThermoFisher) (with primary stain), and 4μg/mL anti-CD16/32 (2.4G2; Bioxcell) to block Fc receptors. The following anti-mouse antibodies were used, with clone and source company listed: CD45 PerCP-Cyanine5.5 (30-F11, BioLegend), CD45 BUV395 (30-F11, BD Horizon), CX3CR1 PE (SA011F11, BioLegend), CX3CR1 PerCP-Cyanine5.5 (SA011F11, BioLegend), CD11b Pacific Blue (M1/70, BioLegend), CD11c BV650 (N418, BioLegend), CD11c Pacific Blue (N418, BioLegend), CD49b Pacific Blue (DX5, BioLegend), CD19 Pacific Blue (6D5, BioLegend), CD44 BV421 (IM7, BioLegend), ST2 Biotin (DIH9, BioLegend), Streptavidin BV510 (BioLegend), PD1 BV605 (29F.1A12, BioLegend), CD3 BV786 (17A2, BioLegend), CD3 APC (17A2, BioLegend), CD4 APC/Cy7 (GK1.5, BioLegend), CD4 PE/Cy7 (GK1.5, BioLegend), CD8 BV650 (53-6.7, BioLegend), CD8 Pacific Blue (53-6.7, BioLegend), CD11a APC (M17/4, eBioscience), CD25 PE/Cy7 (PC61, BioLegend), CD62L BUV737 (MEL-14, BD Horizon), IL-13 PE/Cy7 (eBio13A, Invitrogen), IL-5 APC (TRFK5, BioLegend), GATA3 PerCP-Cyanine5.5 (16E10A23, BioLegend, 1:25). Primary stains were performed for 40 minutes on ice, and secondary staining with Streptavidin was performed for 20 minutes on ice. For intracellular cytokine staining, after isolation from tissue, cells were stimulated with PMA/Ionomycin/GolgiPlug for 4 hours at 37°, 5% CO_2_ prior to any staining. IL-13, IL-5, and GATA3 was stained using the eBioscience Transcription Factor Staining Kit, following the included protocol. Intravascular staining was performed using the CD45 BUV395 antibody according to published protocol (34). Specifically, each mouse was injected with 3μg/150μL retro-orbitally and humanely euthanized 3 minutes post-injection. The BD FACS ARIA II was used for cell sorting, and the Bio-Rad ZE5 Yeti was used for flow cytometry analysis. Data analysis was performed using FlowJo v10 (FlowJo LLC) and Prism.

### Single-cell RNA-sequencing

Single cell suspensions were obtained from livers and spleens as described above. For both tissues, cells were sorted on singlet cells, live cells, CD45^+^ cells, Dump^NEG^ (CD11b^+^, CD11c^+^, CD49b^+^, CD19^+^, CD8^+^) cells, CD3^+^ cells, CD4^+^ cells. Cells were isolated from 2 mice and pooled together, with equal numbers of cells pooled from each animal used. 12,000 cells from each tissue were loaded on a 10X Genomics Next GEM chip and single-cell GEMs were generated on a 10X Chromium Controller. Subsequent steps to generate cDNA and sequencing libraries were performed following 10X Genomics’ protocol. Libraries were pooled and sequenced using Illumina NovaSeq SP 100 cycle as per 10X sequencing recommendations.

The sequenced data were processed using Cell Ranger (version 6.0) to demultiplex the libraries. The reads were aligned to *Mus musculus* mm10 and SCV2 (MN985325.1) genomes to generate count tables that were further analyzed using Seurat (version 4.1.2). Data are displayed as uniform manifold approximation and projection (UMAP). Some plots were generated using the Loupe Cell Browser application from 10X Genomics.

## Results

### CD4^+^ T cells express CX3CR1 primarily in the tissue during both chronic and acute helminth infection

During infection with *Schistosoma mansoni*, parasite eggs become lodged in the liver, inducing immune cell infiltration to form granuloma structures around the eggs. These granuloma structures consist primarily of macrophages, but also eosinophils, and CD4^+^ T cells are needed to produce cytokines that direct and coordinate the response (30). Activated CD4^+^ T cells express CX3CR1 and our previous results suggest a small but significant percentage of CD4^+^ T cells in schistosome granulomas express CX3CR1 (26,28–30). Hence, we used a fate-mapping mouse model in which CX3CR1^CreERT2-IRES-EYFP^ mice were crossed with Rosa26^stop-tdTomato^ reporter mice (hereinafter referred to as CX3CR1^+/CreERT2-eYFP^ R26^+/tdTomato^ mice), which allows for tamoxifen-dependent labeling of CX3CR1^+^ immune cells (12,46). We infected CX3CR1^+/CreERT2-eYFP^ R26^+/tdTomato^ mice with 75 cercariae of *S. mansoni*, and at 7 weeks post-infection, mice were switched to a diet reconstituted with tamoxifen (tamoxifen diet), with uninfected controls placed on tamoxifen diet at the same time. At 8 weeks post-infection, cells from the livers and mesenteric lymph nodes (mesLNs) of both infected mice and uninfected controls were characterized by flow cytometry. We observed a significant increase in the total number of CX3CR1^+^ CD4^+^ T cells in the liver of infected mice as compared to uninfected controls as assessed by antibody staining, as well as a significant increase in total numbers of tdTomato^+^ CD4^+^ T cells in the liver compared to uninfected controls, indicative of active transcription during the 1-week labeling period (Fig1A,B). While there was a slight increase in CX3CR1^+^ CD4^+^ T cell populations in the mesenteric lymph nodes, the magnitude of increase was much less than that which was observed in the liver tissue, suggesting it is primarily in the liver that CX3CR1^+^ CD4^+^ T cells accumulate (Fig1B).

**Figure 1.**
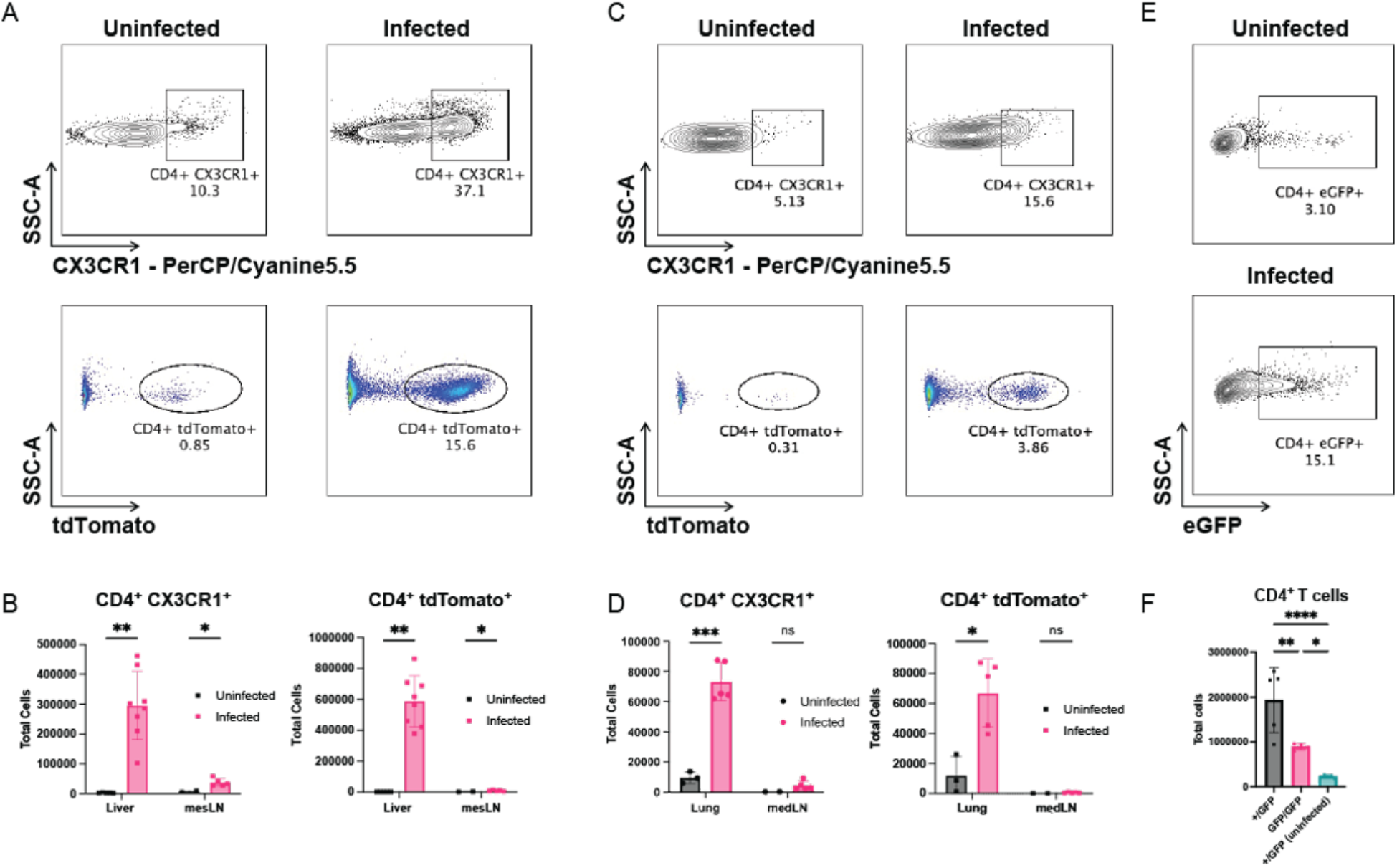
CD4^+^ T cells express CX3CR1 primarily in the tissue during both chronic and acute helminth infection. **(A)** Representative flow cytometry plot of CX3CR1 surface expression and tdTomato labeling on CD4^+^ T cells isolated from the liver of mice infected with *S. mansoni*. **(B)** Quantification of CX3CR1^+^ and tdTomato^+^ CD4^+^ T cells in liver and mesenteric lymph node (mesLN) of mice infected with *S. mansoni*. **(C)** Representative flow cytometry plot of CX3CR1 surface expression and tdTomato labeling on CD4^+^ T cells isolated from the lungs of mice infected with *N. brasiliensis*. **(D)** Quantification of CX3CR1^+^ and tdTomato^+^ CD4^+^ T cells in lungs and mediastinal lymph node (medLN) of mice infected with *N. brasiliensis*. **(E)** Representative flow cytometry plot of eGFP expression in CD4^+^ T cells isolated from the lungs of CX3CR1^eGFP/+^ mice infected with *N. brasiliensis*. **(F)** Quantification of eGFP^+^ CD4^+^ T cells in lungs of CX3CR1^eGFP/+^ and CX3CR1^eGFP/eGFP^ mice infected with *N. brasiliensis*. Cells were gated on singlet cells, live cells, CD45^+^ cells, CD11b^NEG^ cells, CD3^+^ cells, CD8^NEG^, CD4^+^ cells. N = 3-5 mice/ infected group, N = 2-5 mice/ uninfected group, some panels show a combination of two experiments. Error bars represent SEM. Students *t* tests were performed to determine significance. *P<0.05; **P<0.01; ***P<0.001; ****P<0.0001.

While infection with *S. mansoni* is chronic, we also investigated CX3CR1^+^ CD4^+^ T cells with an acute infection in which continuous antigenic stimulation is absent. *Nippostrongylus brasiliensis* is a well-established model of acute helminth infection in mice (31). Following subcutaneous infection, *N. brasiliensis* will travel to the mouse lung in 12-48 hours, causing extensive tissue damage, before they are coughed up and swallowed to gain access to their final niche in the small intestine (31). Infection induces a prominent T_H_2 response in the lung, where CD4^+^ T cells are involved in activating and maintaining innate immune cells involved in the memory response to secondary infection (32). We infected CX3CR1^+/CreERT2-eYFP^ R26^+/tdTomato^ mice subcutaneously with 625 L3 *N. brasiliensis* worms, placing them on tamoxifen diet at the time of infection along with uninfected controls. At day 10 post-infection, we observed a significant increase in total numbers of CD4^+^ T cells expressing CX3CR1 in the lung of infected mice as compared to uninfected controls (Fig1C,D). There was a similar increase in tdTomato^+^ CD4^+^ T cells in the infected lungs compared to uninfected controls. While a small increase in CX3CR1^+^ CD4^+^ T cells was observed in the mediastinal lymph node, it was not statistically significant, and much less than observed in the lung, similar to what was observed during *S. mansoni* infection in the mesenteric lymph node (Fig1D).

We next examined CX3CR1^eGFP^ mice (10), which with homozygosity (CX3CR1^eGFP/eGFP^) has no functional CX3CR1 protein but allows for tracking of cells that would express CX3CR1. We infected both CX3CR1^+/eGFP^ and CX3CR1^eGFP/eGFP^ mice with either 75 cercariae of *S. mansoni* or 625 L3 larvae of *N. brasiliensis* and analyzed either the liver or lungs by flow cytometry, respectively. Uninfected CX3CR1^+/eGFP^ mice in our analysis were used as controls. In the liver of mice infected with *S. mansoni,* we observed a significant increase in the total numbers of eGFP^+^ CD4^+^ T cells as compared to uninfected controls (Supp.Fig1). However, there was no significant difference in the total numbers of CD4 T cells in the infected CX3CR1 deficient mice as compared to the CX3CR1 proficient mice, suggesting CX3CR1 is not needed for CD4 T cell trafficking to the liver during *S. mansoni* infection. In the lungs of mice infected with *N. brasiliensis* we observed a similar increase in the total numbers of eGFP^+^ CD4^+^ T cells (Fig1E) after infection. However, we also observed a lower number of CD4^+^ T cells overall in the lungs of infected CX3CR1 deficient mice as compared to infected CX3CR1 proficient mice (Fig1F), indicating that CX3CR1 plays a role in trafficking to the lungs. Nonetheless, the total number of CD4^+^ T cells was still greater in infected CX3CR1 deficient mice than in the lungs of uninfected mice, suggesting that not all CD4^+^ T cell recruitment to the lung is dependent upon CX3CR1. Other mechanisms are partially responsible for the accumulation CX3CR1 CD4 T cells to the lungs.

Overall, our results demonstrate that CX3CR1^+^ CD4^+^ T cells accumulate primarily at the site of tissue inflammation impacted by helminth infections, instead of the secondary lymphoid organs such as the draining lymph nodes. Additionally, while CX3CR1 does play a role in the accumulation of these CD4 cells in the lungs, it is not required for the accumulation of CX3CR1 CD4^+^ T cells into the liver during *S. mansoni* infection, indicating that there are some tissue environment specific differences in the properties of CX3CR1^+^ CD4^+^ T cells during helminth infections.

### CX3CR1^+^ CD4^+^ T cells have an effector T_H_2 phenotype with greater enrichment of activation markers in the tissue

We next characterized the cellular phenotype of the CX3CR1 CD4 T cells in the tissues. We infected CX3CR1^+/CreERT2-eYFP^ R26^+/tdTomato^ mice with 75 cercariae of *S. mansoni* and placed them on tamoxifen diet at week 7 post-infection with subsequent analysis by flow cytometry at 8 weeks post-infection. Comparing tdTomato^+^ CD4^+^ T cells to tdTomato^-^ CD4^+^ T cells, we found that while tdTomato^-^ CD4^+^ T cells were a mix of activated and naïve cell types, tdTomato^+^ CD4^+^ T cells were uniformly CD44^+^ CD11a^+^ CD62L^-^, indicating an activated T cell phenotype (Fig2A). A higher percentage of tdTomato^+^ cells express ST2, CD25, and PD1 (Fig2B). The higher percentage of cells expressing ST2, the receptor for the alarmin cytokine IL-33, suggests this population is enriched for effector T_H_2 cells (33). CX3CR1^+/CreERT2-eYFP^ R26^+/tdTomato^ mice were also infected with 625 L3 larvae of *N. brasiliensis*, placed on tamoxifen diet at the time of infection, and lungs analyzed at day 10 post-infection. Similar to *S. mansoni* infection, we found that more of the tdTomato^+^ CD4^+^ T cells, as compared to tdTomato^-^ cells, expressed the markers ST2, CD25, and PD1 in the lung following infection (Fig2C).

**Figure 2.**
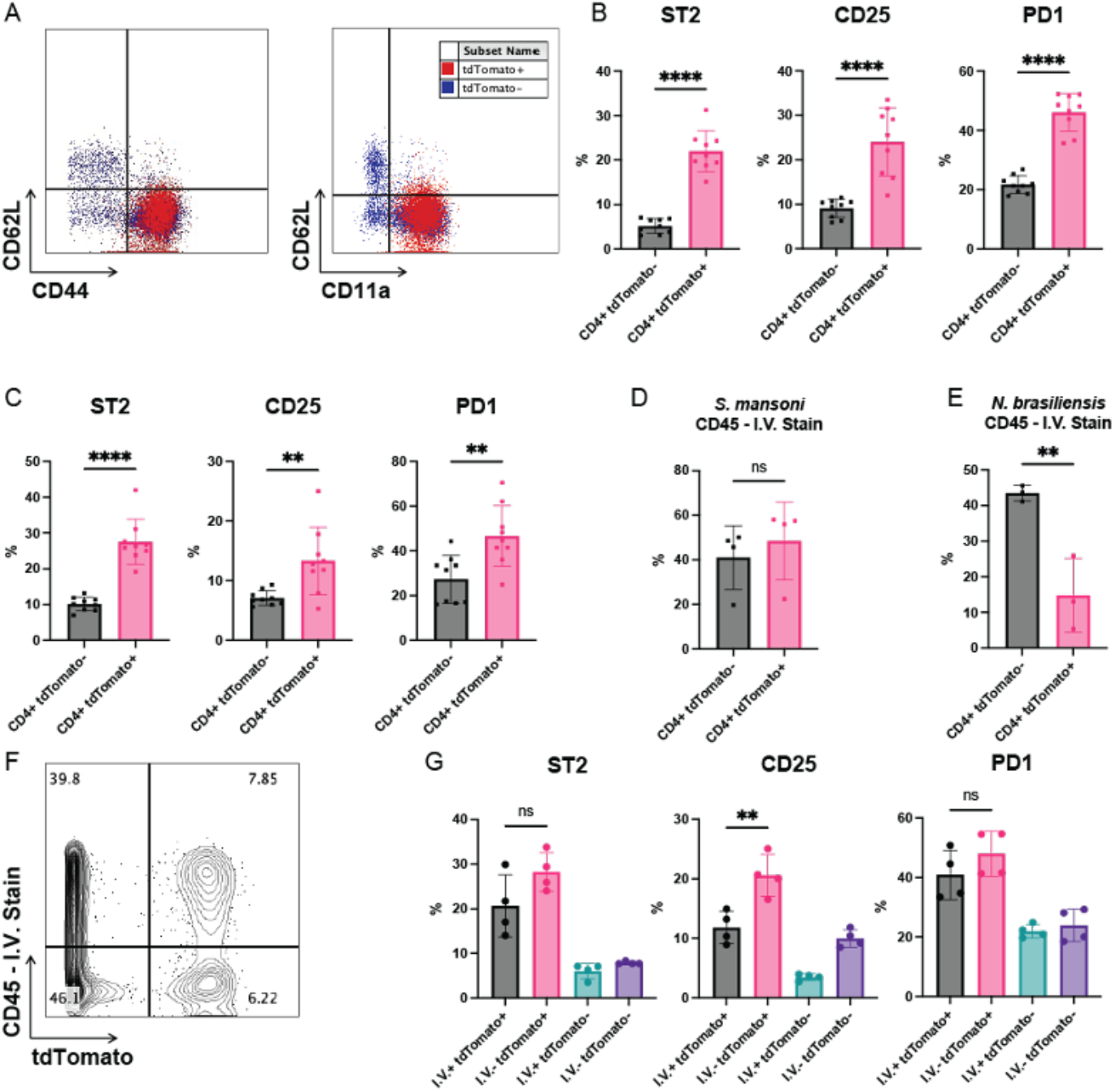
CD4^+^ CX3CR1^+^ cells have an effector T_H_2 phenotype with greater enrichment of activation markers in the tissue. **(A)** Representative overlayed flow cytometry plots of CD44, CD11a, and CD62L expression on both tdTomato^+^ and tdTomato^-^ CD4^+^ T cells isolated from the liver of mice infected with *S. mansoni*. **(B)** Percentage of tdTomato^+^ and tdTomato^-^ CD4^+^ T cells in liver of mice infected with *S. mansoni* that express ST2, CD25, and PD1. **(C)** Percentage of tdTomato^+^ and tdTomato^-^ CD4^+^ T cells in lungs of mice infected with *N. brasiliensis* that express ST2, CD25, and PD1. **(D)** Percentage of tdTomato^+^ and tdTomato^-^ CD4^+^ T cells positive for the anti-CD45 intravascular stain in liver of mice infected with *S. mansoni* and **(E)** lungs of mice infected with *N. brasiliensis*. **(F)** Representative flow cytometry plot of anti-CD45 staining versus tdTomato expression in CD4^+^ T cells in liver of mice infected with *S. mansoni.* **(G)** Percentage of tdTomato^+^ and tdTomato^-^,CD45^+^ and CD45^-^ CD4 T cells in liver of mice infected with *S. mansoni* that express ST2, CD25, and PD1. Cells were gated on singlet cells, live cells, CD11b^NEG^ cells, CD3^+^ cells, CD8^NEG^, CD4^+^ cells. N = 3-5 mice/group, some panels show a combination of two experiments. Error bars represent SEM. Students *t* tests were performed to determine significance. *P<0.05; **P<0.01; ***P<0.001; ****P<0.0001.

We next performed intravascular labeling of immune cells with anti-CD45 fluorescent antibody, labeling circulating immune cells, leaving cells in the parenchyma unlabeled (34). In the livers of mice infected with *S. mansoni* we observed an equal percentage of tdTomato^+^ and tdTomato^-^ CD4^+^ T cells positive for the intravascular CD45 stain, suggesting equal levels of tissue infiltration for the two populations (Fig2D). However, in the lungs of mice infected with *N. brasiliensis*, we observed a significantly lower percentage tdTomato^+^ CD4^+^ T cell population that stained positive for CD45 compared to the tdTomato^-^ population, suggesting the tdTomato^+^ cells preferentially transit into the parenchyma (Fig2E). We also compared surface marker expression between intravascular^+^ (I.V.^+^) and intravascular^-^ (I.V.^-^) tdTomato^+^ CD4^+^ T cells in the liver after *S. mansoni* infection and observed more I.V.^-^ tdTomato^+^ cells expressing CD25, as well as the same trends for ST2 and PD1, compared to I.V.^+^ tdTomato^+^ cells, suggesting a greater enrichment of activation markers once the cells have transited into the tissue (Fig2F,G).

Overall, these results indicate that the fate mapped CD4^+^ cells that have expressed CX3CR1 in the tissues of helminth infected mice have primarily the surface phenotype of effector T_H_2 cells. However, while they preferentially accumulated in the parenchyma of the lungs during *N. brasiliensis* infection, in the liver these cells are equally distributed between sinusoids and tissues, indicating tissue specific differences in migration.

### CX3CR1^+^ CD4^+^ T cells produce less IL-4 despite expressing GATA3

We used the 4get mice (for ‘IL-4/GFP-enhanced transcript’) (35) to examine IL-4 expression in CX3CR1^+^ CD4^+^ T cells after *S. mansoni* or *N. brasiliensis* infection and analyzed either the liver or the lungs, respectively, at either 8 weeks post-infection for the liver, and 10 days post-infection for the lung. As expected, in both tissues isolated from infected 4get mice, there was a significant increase in eGFP^+^ CD4^+^ T cells (Supp.Fig2C,D). Surprisingly, while there are more CX3CR1^+^ CD4^+^ T cells upon infection, IL-4 eGFP expression is primarily found in the CX3CR1^-^ CD4^+^ T cells (Fig3E-H). This suggests that CX3CR1^+^ CD4^+^ T cells are not the main source of IL-4 during helminth infections. We next performed intracellular staining on cells from the liver or lungs to assess cytokine production and transcription factor expression. In both types of helminth infections of mice, there was an increase of GATA3^+^ CD4^+^ T cells (Supp.FigA,B).

**Figure 3.**
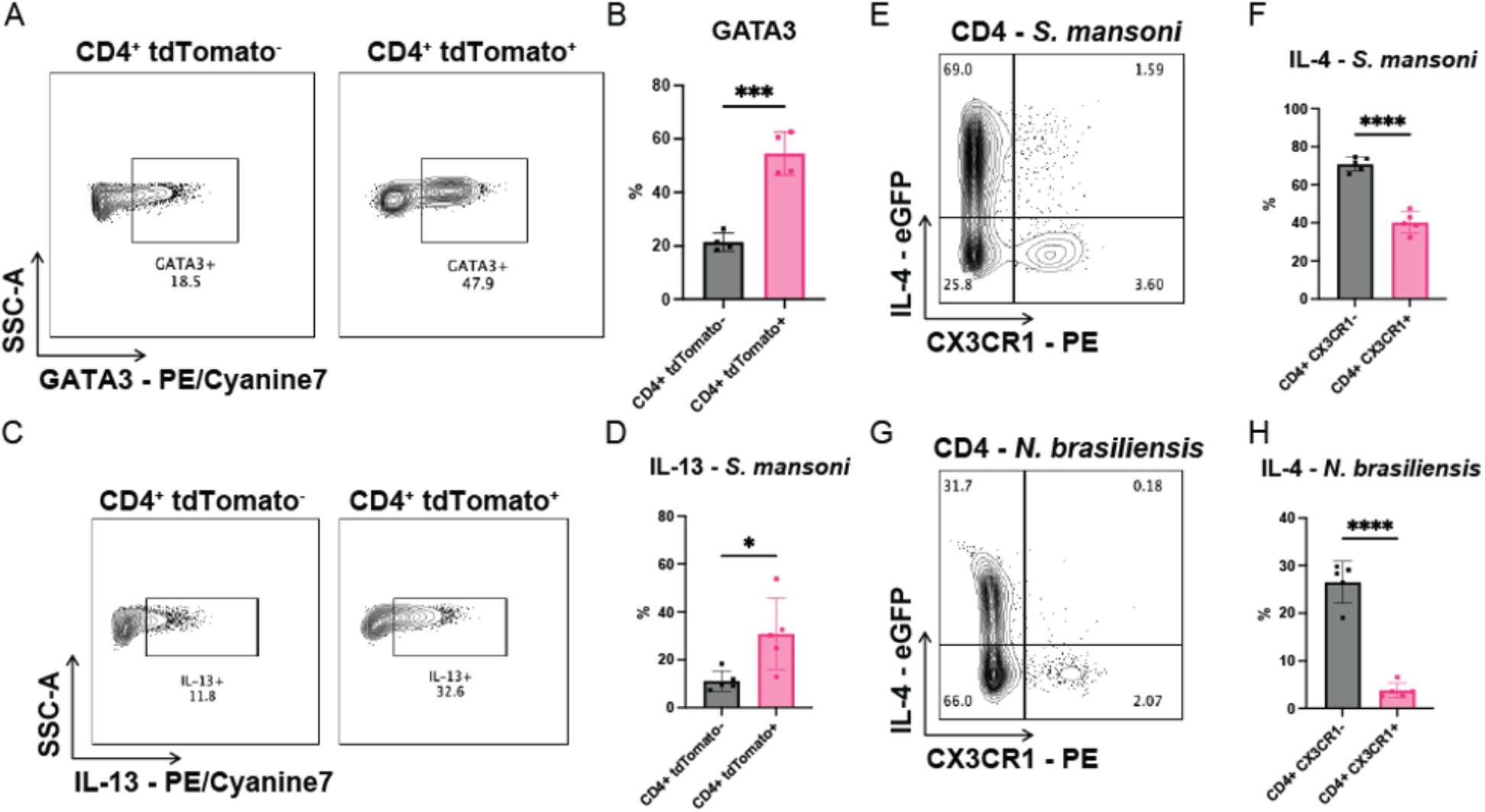
CD4^+^ tdTomato^+^ cells express GATA3 and are enriched for IL-13. **(A)** Representative flow cytometry plots of GATA3 expression on tdTomato^+^ and tdTomato^-^ CD4^+^ T cells isolated from the liver of mice infected with *S. mansoni*. **(B)** Percentage of tdTomato^+^ and tdTomato^-^ CD4^+^ T cells in liver of mice infected with *S. mansoni* that express GATA3. **(C)** Representative flow cytometry plots of IL-13 expression in tdTomato^+^ and tdTomato^-^ CD4^+^ T cells in mice infected with *S. mansoni.* **(D)** quantification of percentage of IL- 13 expression in tdTomato^+^ and tdTomato^-^ CD4^+^ T cells in mice infected with *S. mansoni.*. **(E)** Representative flow cytometry plot of IL-4 expression versus CX3CR1 surface staining in CD4^+^ T cells in liver of 4get mice infected with *S. mansoni.* **(F)** Percentage of CX3CR1^+^ and CX3CR1^-^ CD4 T cells in 4get mice infected with *S. mansoni* **(G)** Representative flow cytometry plot of IL- 4 expression versus CX3CR1 surface staining in CD4^+^ T cells in lungs of 4get mice infected with *N. brasiliensis.* (H) Percentage of CX3CR1^+^ and CX3CR1^-^ CD4 T cells in 4get mice infected with *N. brasiliensis*. Cells were gated on singlet cells, live cells, CD45^+^ cells, CD11b^NEG^ cells, CD3^+^ cells, CD8^NEG^, CD4^+^ cells. N = 3-5 mice/group. Error bars represent SEM. Students *t* tests were performed to determine significance. *P<0.05; **P<0.01; ***P<0.001; ****P<0.0001.

In the livers of mice infected with *S. mansoni*, we observed about 20% of tdTomato^-^ CD4^+^ T cells expressing GATA3, while about 50% of tdTomato^+^ CD4^+^ T cells expressed GATA3, consistent with this population being enriched for the T_H_2 phenotype (Fig3A,B). More tdTomato^+^ CD4^+^ T cells were also observed to produce IL-13 compared to tdTomato^-^ CD4^+^ T cells (Fig3C,D). Similar results were seen when comparing expression of IL-5, although expression was lower overall in both populations, and the difference between the two was not statistically significant. (Supp.Fig2E,F). These results indicate that CX3CR1^+^ CD4^+^ T cells are producers of effector T_H_2 cytokines IL-5 and IL-13, but not IL-4, despite GATA3 expression.

### tdTomato^+^ CD4^+^ T cells that retain CX3CR1 expression persist longer in the peripheral tissues

By utilizing the fate-mapping CX3CR1^+/CreERT2-eYFP^ R26^+/tdTomato^ mouse, we can track the dynamics of CD4^+^ T cells that have expressed CX3CR1. We infected CX3CR1^+/CreERT2-eYFP^ R26^+/tdTomato^ mice with *S. mansoni* and placed them on tamoxifen diet at week 7 post-infection.

One cohort of mice was analyzed at week 8 post-infection, and the second cohort was taken off tamoxifen diet at week 8 but left until week 12 post-infection for analysis. There are significantly fewer tdTomato^+^ CD4^+^ T cells at the later timepoint, indicating that the cells that were initially labeled had died off, although there was still more than in uninfected controls (Fig4A). At both timepoints, we also assessed CX3CR1 expression by antibody staining on tdTomato^+^ CD4^+^ T cells. At week 8 post-infection, about 70% of tdTomato^+^ CD4^+^ T cells were CX3CR1^+^, indicating a small but significant percentage of cells downregulated CX3CR1 after being labeled (Fig4C,D). At week 12 post-infection, the percentage of tdTomato^+^ CD4^+^ T cells expressing CX3CR1 increased to over 90%, despite lower total numbers of cells overall, suggesting those cells that maintained CX3CR1 expression were better able to persist in the liver (Fig4C,D). We similarly assessed ST2 expression at week 8 and week 12 timepoints as a proxy for T_H_2 effector function. At week 8 post-infection, about 20% of tdTomato^+^ CD4^+^ T cells expressed ST2, while at week 12 post-infection, about 50% of tdTomato^+^ CD4^+^ T cells that persisted expressed ST2 (Fig4E).

**Figure 4.**
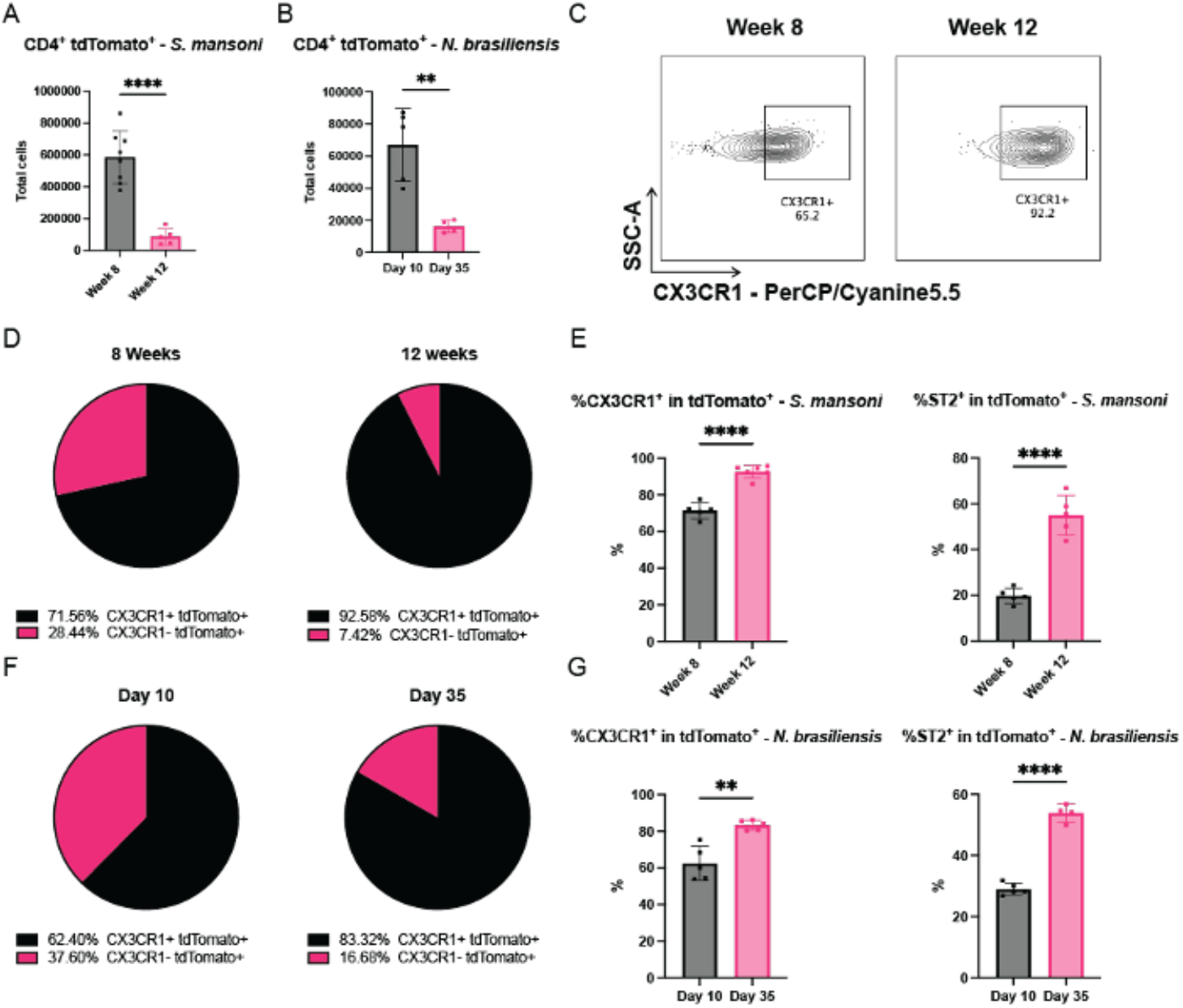
CD4^+^ CX3CR1^+^ cells persist longer in the peripheral tissues while maintaining ST2 expression. **(A)** Quantification of total tdTomato^+^ CD4^+^ T cells isolated from the liver of mice infected with *S. mansoni* at 8 and 12 weeks post-infection and **(B)** from lungs of *N. brasiliensis* at 10 and 35 days post-infection **(C)** Representative flow cytometry plot of CX3CR1 expression on tdTomato^+^ CD4^+^ T cells in liver of mice infected with *S. mansoni* at both 8 and 12 weeks post-infection **(D)** Pie graphs of average CX3CR1 expression on tdTomato^+^ CD4^+^ T cells mice infected with *S. mansoni* at 8 and 12 weeks post-infection **(E)** Quantification of CX3CR1 and ST2 expression on tdTomato^+^ CD4^+^ T cells in liver of *S. mansoni* infected mice at 8 and 12 weeks post-infection. **(F)** Pie graphs of average CX3CR1 expression on tdTomato^+^ CD4^+^ T cells mice infected with *N. brasiliensis* at 10 and 35 weeks post-infection **(G)** Quantification of CX3CR1 and ST2 expression on tdTomato^+^ CD4^+^ T cells in lungs of *N. brasiliensis* infected mice at 10 and 35 days post-infection. Cells were gated on singlet cells, live cells, CD45^+^ cells, CD11b^NEG^ cells, CD3^+^ cells, CD8^NEG^, CD4^+^ cells. N = 3-5 mice/group. Error bars represent SEM. Students *t* tests were performed to determine significance. *P<0.05; **P<0.01; ***P<0.001; ****P<0.0001.

To compare with an acute infection model, we next infected CX3CR1^+/CreERT2-eYFP^ R26^+/tdTomato^ mice with L3 larvae of *N. brasiliensis* and placed them on tamoxifen diet at the time of infection. At day 10 post-infection, we analyzed the lungs of one cohort, while taking the other cohort off tamoxifen diet, which was analyzed at day 35 post-infection. As with the liver, there was a significant decrease in tdTomato^+^ CD4^+^ T cells from day 10 to day 35, although the numbers at day 35 remained higher than in that of uninfected controls (Fig4B). At day 10 post-infection, about 60% of tdTomato^+^ CD4^+^ T cells expressed CX3CR1 on their surface, while at day 35 over 80% of tdTomato^+^ CD4^+^ T cells expressed CX3CR1 (Fig4F,G). Similarly, we saw more ST2 expression within the tdTomato^+^ CD4^+^ T cell population at day 35 post-infection (Fig4G). Overall, these data suggests that in both chronic and acute helminth infection, the CD4^+^ T cells that maintain CX3CR1 surface expression as well as the effector T_H_2 phenotype persist for longer in the afflicted tissue.

### CX3CR1-fate-mapped CD4^+^ T cells have distinct phenotypes in the spleen compared to peripheral tissues

While we found the accumulation of CX3CR1^+^ CD4^+^ T cells primarily in the peripheral tissue affected by parasites, we also observed a slight but significant increase in tdTomato^+^ CD4^+^ T cells in the spleens of CX3CR1^+/CreERT2-eYFP^ R26^+/tdTomato^ mice infected with either *S. mansoni* or *N. brasiliensis* and placed on tamoxifen diet as described above (Fig5A,B). To further assess splenic tdTomato^+^ CD4^+^ T cells during infection, we investigated CX3CR1^+/CreERT2-eYFP^ R26^+/tdTomato^ mice infected with *S. mansoni* at both 8- and 12-weeks post-infection, after labeling with tamoxifen diet from week 7 to week 8. At week 8 post-infection, tdTomato^+^ CD4^+^ T cells in the spleen appeared with a similar phenotype to tdTomato^+^ CD4^+^ T cells found in the liver, with only minor differences (Fig5C, Supp.Fig3A). Cells in the spleen were CD44^+^ and mostly CD11a^+^, with a small number of cells remaining CD11a^-^. However, some tdTomato^+^ CD4^+^ T cells in the spleen were both CD44^+^ and CD62L^+^, indicative of a central memory (T_CM_) – like phenotype. While there are fewer tdTomato^+^ CD4^+^ T cells in the spleen at week 12 post-infection, the total numbers at week 12 were still higher than those in the uninfected controls, suggesting that labeled cells can persist in the spleen (Fig5D).

**Figure 5.**
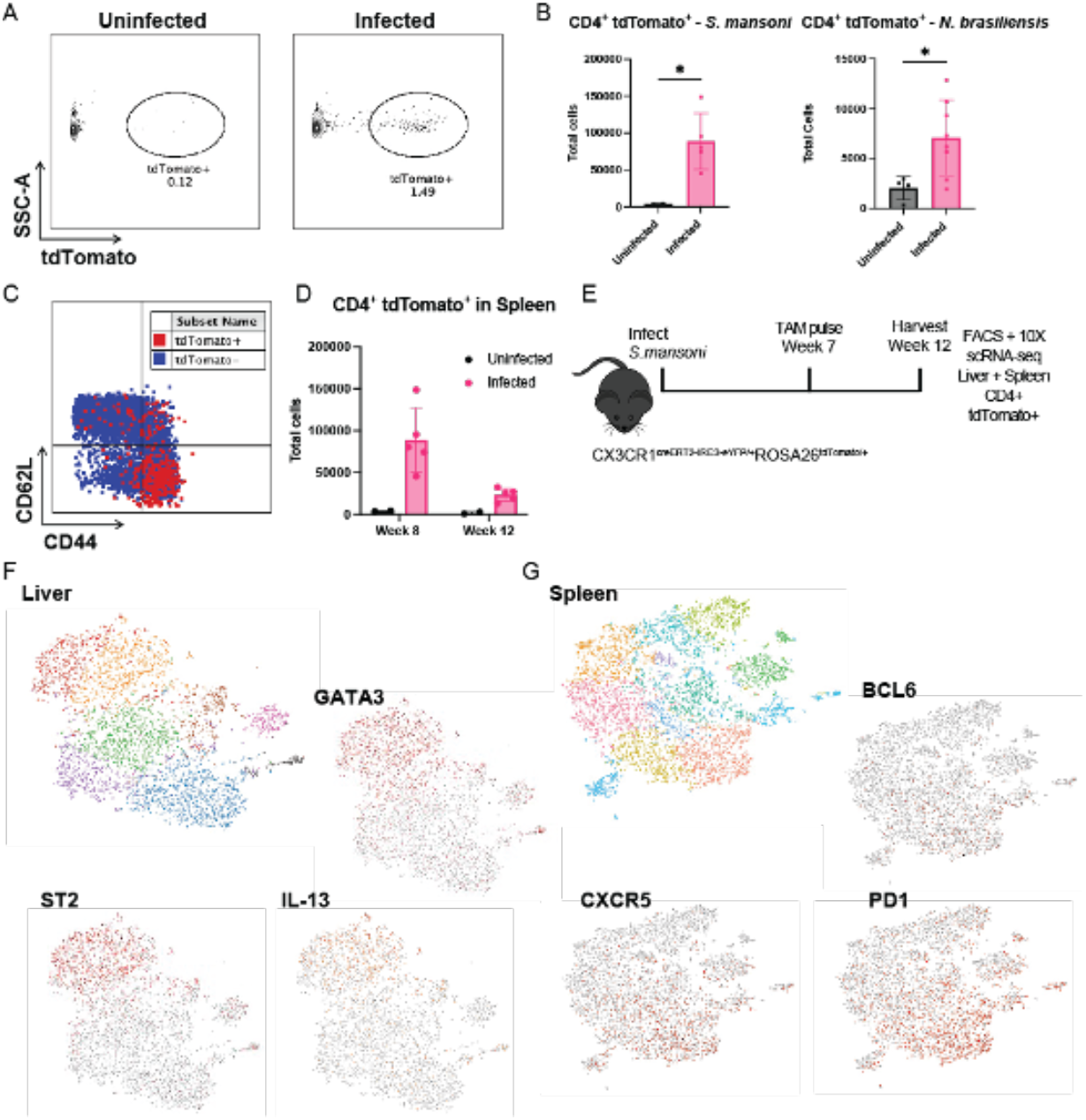
CD4^+^ CX3CR1^+^ fate-mapped cells have distinct phenotypes in the spleen compared to peripheral tissues. **(A)** Representative flow cytometry plot of tdTomato^+^ CD4^+^ T cells in spleen of *S. mansoni* infected mice **(B)** Quantification of tdTomato^+^ CD4^+^ T cells in spleen of *S. mansoni* and *N. brasiliensis* infected mice **(C)** Overlayed representative flow cytometry plot of CD44 and CD62L expression on both tdTomato^+^ and tdTomato^-^ CD4^+^ T cells isolated from the spleen of mice infected with *S. mansoni* at 8 weeks post-infection **(D)** Quantification of tdTomato^+^ CD4^+^ T cells in spleen of *S. mansoni* at 8 and 12 weeks post- infection. **(E)** Experimental design of experiment to label and sort out tdTomato^+^ CD4^+^ T cells from both the liver and spleen **(F)** UMAP clustering of tdTomato^+^ CD4^+^ T cells from the liver of mice infected with *S. mansoni* with selected genes highlighted. **(G)** UMAP clustering of tdTomato^+^ CD4^+^ T cells from the spleen of mice infected with *S. mansoni* with selected genes highlighted. For analysis, cells were gated on singlet cells, live cells, CD45^+^ cells, CD11b^NEG^ cells, CD3^+^ cells, CD8^NEG^, CD4^+^ cells. For sorting/scRNA-seq, cells were sorted as described in methods. N = 3-5 mice/infected group, N = 2-5 mice/uninfected group, some panels show a combination of two experiments. Error bars represent SEM. Students *t* tests were performed to determine significance. *P<0.05; **P<0.01; ***P<0.001; ****P<0.0001.

Since the surface phenotype of some splenic tdTomato^+^ CD4^+^ T cells was different from the liver tdTomato^+^ CD4^+^ T cells, we next determined gene expression differences between the populations of tdTomato^+^ CD4^+^ T cells in both the spleen and the liver that occur during *S. mansoni* infection by scRNA-seq. At week 7 post-infection we administered tamoxifen to all mice via oral gavage, to create a single labeling event. At week 12 post-infection, we FACS isolated tdTomato^+^ CD4^+^ T cells from both the liver and the spleen and submitted 12,000 cells from each population for single-cell RNA-sequencing (Fig5E). We first visualized each population separately with UMAP hierarchical clustering, and defined multiple clusters based on gene expression. In the liver, there appeared to be two major populations: one displaying gene expression consistent with a T_H_2 phenotype, with high expression of GATA3, IL-13, and ST2, and the other consistent with a T_H_1 phenotype, with high expression of IFNγ, Gzmk, and CXCR3 (Fig5F, Supp.Fig3B). In the spleen there was also considerable heterogeneity, with distinct IFNγ^+^ and GATA3^+^ clusters, but also high expression of LAG3, which is often associated with a T cell state known as exhaustion in which activated cells are less responsive (Fig5G, Supp.Fig3C) (36). Notably, a significant number of cells in the spleen were positive for BCL6, CXCR5, and PD1, all markers of the follicular T helper (T_FH_) subset of CD4 T cells (Fig5G) (37). Our analysis revealed a considerable amount of heterogeneity in tdTomato^+^ CD4^+^ T cell populations, both in the liver and the spleen, suggesting that CX3CR1 does not strictly define one subset of CD4^+^ T cells, but instead is a general marker of activation for a number of different CD4^+^ T cell populations. Over time, CD4^+^ cells expressing CX3CR1 can go on to adopt a multitude of different phenotypic profiles, rather than follow a specific differentiation path.

### CX3CR1-specific deletion of BCL6 reduces accumulation of CX3CR1^+^ CD4^+^ T cells in the spleen during *N. brasiliensis* infection

UMAP visualization on the scRNA-seq datasets helped identify distinguishing transcription factors between tdTomato^+^ CD4^+^ T cells isolated from either the liver or the spleen (Fig6A). Cells from the liver were characterized by high expression of the transcription factor ID2, while cells from the spleen expressed high levels of TCF7 and TOX (Fig6B). Notably, we observed BCL6 expression almost exclusively in the spleen, suggesting that some of the tdTomato^+^ CD4+ T cells in the spleen are T_FH_ cells. We also confirmed in our flow cytometry data from mice infected either with *S. mansoni* or *N. brasiliensis* that a portion of tdTomato^+^ CD4^+^ T cells were CD44^+^ PD1^+^ in both infections, akin to a T_FH_ phenotype (Fig6C).

**Figure 6.**
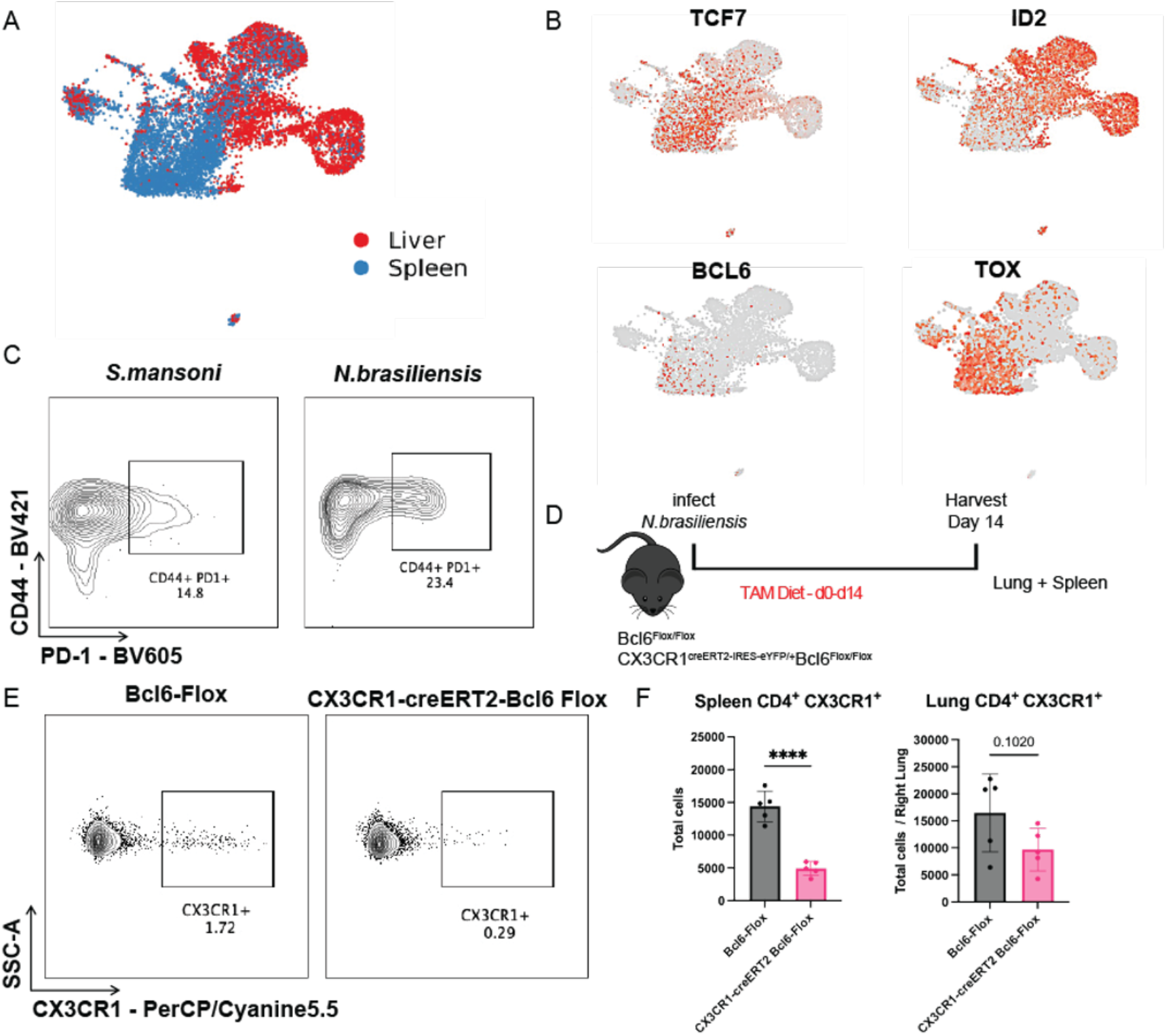
CX3CR1-specific deletion of BCL6 reduces accumulation of CD4^+^ CX3CR1^+^ cells in the spleen during *N. brasiliensis* infection. **(A)** Combined UMAP clustering of tdTomato^+^ CD4^+^ T cells from both the spleen and liver of mice infected with *S. mansoni* (B) Expression patterns of transcription factors in combined UMAP clustering of cells from both liver and spleen **(C)** Expression of CD44 and PD1 on tdTomato^+^ CD4^+^ T cells in spleen of *S. mansoni* and *N. brasiliensis* infected mice **(D)** Experimental design of experiment to conditionally delete Bcl6 in CX3CR1^+^ cells **(E)** representative flow cytometry plots of CX3CR1 expression in the spleen **(F)** Quantification of CX3CR1^+^ CD4^+^ T cells in the spleen and lungs of Bcl6^Flox^ and CX3CR1^CreERT2^ Bcl6^Flox^ mice infected with *N. brasiliensis* and. For analysis, cells were gated on singlet cells, live cells, CD45^+^ cells, CD11b^NEG^ cells, CD3^+^ cells, CD8^NEG^, CD4^+^ cells. For sorting/scRNA-seq, cells were sorted as described in methods. N = 5 mice/group. Error bars represent SEM. Students *t* tests were performed to determine significance. *P<0.05; **P<0.01; ***P<0.001; ****P<0.0001.

The expression of BCL6 in the tdTomato^+^ CD4^+^ T cell population was intriguing as it is part of a complex transcriptional network and may play a role in differentiation and survival of T_FH_ cells as well as other CD4^+^ T cell populations (38). We generated a conditional knockout mouse model by crossing the CX3CR1^CreERT2-IRES-eYFP^ mice to mice in which the BCL6 gene has been flanked by loxP sequences, to generate a mouse with the genotype CX3CR1^+/CreERT2-IRES-eYFP^ Bcl6^Flox/Flox^, hereinafter referred to as CX3CR1^CreERT2^ Bcl6^Flox^ mice. Exposure to tamoxifen results in the deletion of BCL6 from cells expressing CX3CR1 at the time. We used littermate controls with the genotype CX3CR1^+/+^ Bcl6^Flox/Flox^, hereinafter referred to as Bcl6^Flox^. We infected both CX3CR1^CreERT2^ Bcl6^Flox^ and Bcl6^Flox^ mice with *N. brasiliensis*, placing both groups on tamoxifen at the time of infection (Fig6D). We analyzed the lungs and spleens of infected mice at day 14 to ensure deletion of BCL6 throughout the course of infection (Fig3.12A).

Compared to Bcl6^Flox^ controls, CX3CR1^CreERT2^ Bcl6^Flox^ mice had significantly lower numbers of CX3CR1^+^ CD4^+^ T cells present in the spleen, suggesting that BCL6 is important for maintaining this population (Fig6E,F). We also found slightly reduced numbers of CX3CR1^+^ CD4^+^ T cells in the lungs of CX3CR1^CreERT2^ Bcl6^Flox^ mice compared to controls, although it was not statistically significant (Fig6F). However, there was not a significant reduction in the number of PD1^+^ CXCR5^+^ T_FH_ cells (Supp.Fig4A). Overall, our results show that during *N. brasiliensis* infection, BCL6 plays an important role to maintain populations of CX3CR1^+^ CD4 T cells in the spleen, but this does not affect the overall abundance of T_FH_ cells in the spleen. We next infected both CX3CR1^CreERT2^ Bcl6^Flox^ and Bcl6^Flox^ mice with *S. mansoni*, placing both groups on tamoxifen at week 7 post-infection, up until the point of analysis at week 12 post-infection (Supp.Fig4B).

Unlike during *N. brasiliensis* infection, we did not observe a significant difference in total numbers of CX3CR1^+^ CD4^+^ T cells in the spleen (Supp.Fig4B). However, we did find reduced numbers of CX3CR1^+^ CD4^+^ T cells in the livers of CX3CR1^CreERT2^ Bcl6^Flox^ mice compared to controls, indicating a role for BCL6 in maintaining this population in the affected tissue (Supp.Fig4C). Overall, our results show that during *S. mansoni* infection, BCL6 is needed to maintain populations of CX3CR1^+^ CD4^+^ T cells in the liver, suggesting that BCL6 can play a role in maintaining CX3CR1^+^ CD4^+^ T cells in the tissue as well as the spleen depending on whether the helminth infection is chronic or acute.

## Discussion

In this study, we used a CX3CR1 fate-mapping mouse model to characterize CX3CR1^+^ CD4^+^ T cells during models of both acute (*N. brasiliensis*) and chronic (*S. mansoni*) helminth infection and find that they primarily accumulate as T_H_2 effector cells in peripheral tissues but can also be found in the spleen with a distinct phenotype. By combining single-cell RNA-sequencing with fate mapping CX3CR1^+^ CD4^+^ cells, we found a population of fate-mapped splenic CD4^+^ T cells that are TCF-1^+^, TOX^+^, and BCL6^+^ during *Nippostrongylus brasiliensis* infection. Conditional deletion of BCL6 in this population in CX3CR1^CreERT2^ Bcl6^Flox/Flox^ mice treated with tamoxifen indicates a role for BCL6 in maintaining this population of CD4^+^ T cells.

The expression of CX3CR1 as a marker on T cells during immune responses has so far focused primarily on either CD8^+^ T cells (23–25), or cytotoxic T_H_1 CD4^+^ T cells (26,27). Here we follow the dynamics of CX3CR1^+^ CD4^+^ T cells during type 2 inflammatory responses, resultant from both chronic and acute helminth infections. During chronic viral infections, CX3CR1^+^ CD8^+^ T cells are recent progeny of stem-like TCF-1^+^ CD8^+^ T cell cells and checkpoint blockade of the PD1 pathway can increase the abundance of these cells, which may contribute to viral control and tumor immunity (51). Similarly, we find here that accumulation of CX3CR1^+^ CD4^+^ cells during *N. brasiliensis* and *S. mansoni* is primarily at the site of inflammation, in this case either the lung or the liver. We found that fate-mapped tdTomato^+^ CD4^+^ T cells in the tissue were uniformly CD44^+^ CD11a^+^ CD62L^-^, a phenotype consistent with T cell activation. Furthermore, tdTomato^+^ CD4^+^ T cells were enriched in expression of ST2, IL- 13, and GATA3, suggesting differentiation into a T_H_2 phenotype. These results are consistent with other studies that have shown CX3CR1 expression on CD4^+^ T cells is mostly a condition of proximity to inflammation, and an absence of expression on naïve CD4 T cells (17,26,28).

While the cellular dynamics and phenotype of CX3CR1^+^CD4^+^ cells were similar in both acute (*N. brasiliensis*) and chronic (*S. mansoni*) infection models, there were some important discrepancies between the two. During acute *N. brasiliensis* infection, CX3CR1 plays an important role in the accumulation of the CD4^+^ T cells in the lungs, and CX3CR1 deficiency resulted in lower numbers of total CD4^+^ T cells in the lung, although it was still higher than in uninfected controls. In contrast, during *S. mansoni* infection, CX3CR1 deficiency had no effect on total numbers of CD4^+^ T cells present in the liver at the peak of infection. We speculate that this may be because *S. mansoni* is a chronic infection in which immune cells are being drawn into the site of inflammation over an extended period, whereas in *N. brasiliensis* infection, antigenic stimulus in the lung occurs only while the worms are present for a couple of days at most, and thus there might be less redundancy and intensity in factors driving T cell recruitment, causing CX3CR1 deficiency to have a greater effect. Additionally, we observed differences in the intravascular staining of tdTomato^+^ CD4^+^ T cells between the two infections. During *N. brasiliensis* infection, most tdTomato^+^ CD4^+^ T cells in the lung were negative for the intravascular stain, while in *S. mansoni* infection this population was evenly split between the parenchyma and the vasculature. This could be due to continuous recruitment of CX3CR1^+^ CD4^+^ T cells to the liver during *S. mansoni* infection, thus diluting the percentage of these cells that have crossed into the tissue. While CX3CR1 can be involved in immune cell trafficking, this function of CX3CR1 on CD4 T cells is likely to be context specific. For example, one study found in a model of LCMV13 infection CX3CR1 to be dispensable for T cell trafficking to the spleen, lung, and liver, while others have found it necessary in models of *Helicobacter pylori* infection and allergic asthma (26,28,39). Our own analysis on two different models of helminth infection affecting two different tissues supports this view that CX3CR1 is involved in CD4^+^ T cell trafficking in some instances but not others.

CX3CR1 is considered a surface marker of T cell activation, with initial studies suggesting it was primarily a marker of T_H_1 CD4^+^ T cells, but it is also expressed on T_H_2 cells in a model of allergic asthma in which CX3CR1^+^ CD4^+^ T cells drove pathology (28,40). In a model of atopic dermatitis, both T_H_1 and T_H_2 CD4^+^ T cells could express CX3CR1 (29). Hence, expression of CX3CR1 could be indicative of an activation state, rather than a differentiation state, with considerable heterogeneity of functions. Our results here support the view that CX3CR1^+^ CD4^+^ T cells can adopt a variety of phenotypes over time. ScRNA-seq analysis of tdTomato^+^ CD4^+^ T cells isolated from *S. mansoni* infected mice, 5 weeks after selective labeling, revealed considerable heterogeneity between populations isolated from either the liver or the spleen. Cells in the liver were defined by the transcription factor ID2, which preferentially mark effector CD4^+^ T cells, while cells in the spleen predominantly expressed TCF7, a transcription factor associated with less differentiated T cells (41–44). However, even within these two populations there was considerable heterogeneity, with the liver containing both T_H_1 and T_H_2 CD4^+^ T cell phenotypes while the spleen had a mix of cell phenotypes, including a T_FH_ population. We initially hypothesized that accumulated CX3CR1^+^ CD4^+^ T cells in the spleen are precursors to T_FH_ cells during helminth infections, however the conditional deletion of BCL6 in CX3CR1^+^ cells did not significantly reduce the frequency of T_FH_ cells during infection, although it diminishes the overall number of CX3CR1^+^ CD4^+^ cells. Hence, CX3CR1 is probably expressed in a subset of T_FH_ cells during infection, but is not a critical marker for all the T_FH_ cells activated during infection. Additionally, our experiments with the 4get mice indicates that CX3CR1^+^ CD4^+^ cells do not express IL-4 transcripts, which is also an important indicator of T_FH_ cells.

Utilizing the CX3CR1^+/CreERT2-eYFP^ R26^+/tdTomato^ fate-mapping system allowed for us to track the dynamics of CX3CR1-expressing cells overtime, regardless of whether they downregulate CX3CR1. An important observation was that tdTomato^+^ CD4^+^ T cells that maintained CX3CR1 expression had a survival advantage over those that did not, evidenced by the enrichment of CX3CR1 in tdTomato^+^ CD4^+^ T cells at later timepoints. While we do not know the exact mechanism, there is evidence in the literature to suggest that the downstream signaling from CX3CR1 upregulates anti-apoptotic factors such as BCL2 that could improve cellular survival (21,22). Furthermore, studies of CD8^+^ T cells suggest CX3CR1 marks terminal effector memory T cells (terminal-T_EM_) that persist after infection (25). While no comparative study has been done on CD4^+^ T cells, it is possible that in our own experiments CX3CR1 is marking a subset of memory CD4^+^ T cells that persist in the tissue. Another notable result from our study was that conditional deletion of BCL6 from CX3CR1 expressing cells resulted in lower levels of CX3CR1^+^ CD4^+^ T cells in the spleen during *N. brasiliensis* infection and the liver during *S. mansoni* infection. While we predicted there might be an effect on this population in the spleen, we were surprised to find a reduction of CX3CR1^+^ CD4^+^ T cells in the liver during chronic infection. While BCL6 is well known as the lineage defining transcription factor of T_FH_ cells, it has been shown to be part of a much broader transcriptional network directing CD4^+^ T cell behavior (38,45). It is possible that by deleting BCL6, we disrupted this broader network in some unknown way that prevented CX3CR1^+^ CD4^+^ T cell accumulation. An important limitation of our studies is that the exact function of the CX3CR1^+^ CD4^+^ during these helminth infections is still unclear. Our efforts to specifically deplete this population using a Diptheria Toxin system was unsuccessful. Additionally, CX3CR1 appears to be a marker that is expressed in a heterogenous fashion and hence may not mark specifically functionality during a response.

While CX3CR1^+^ CD4^+^ T cells could be an important source of IL-5 and IL-13 effector T_H_2 cytokines, there are still abundant effector T_H_2 cells producing these cytokines that are not CX3CR1^+^. Hence, it is still unclear if expression of CX3CR1 is simply an activation marker for a subset of T_H_2 cells, or whether this population of CD4^+^ T cells has a specific immune function.

Helminth infections continue to be a major public health threat, with an urgent need for an effective vaccine against both hookworm and schistosome infection. Development of an effective vaccine is contingent upon a detailed yet broad understanding of the immune response to these pathogens. Our study contributes to that effort by characterizing a population of CX3CR1-expressing CD4^+^ T cells in models of both acute and chronic infection. We found these cells to be uniformly activated in both instances, and maintenance of CX3CR1 expression providing a survival advantage to T_H_2 cell persisting in the tissue. However, more work is needed to define the exact role of these cells during infection, and how they might be contributing to immunity in the context of helminth infections.

## Acknowledgements

We acknowledge help from the NYU Langone’s Genome Technology Center for performing all the RNA sequencing. Cell sorting/flow cytometry technologies were provided by NYU Langone’s Cytometry and Cell Sorting Laboratory. The NYU Langone Health Genome Technology Center and the Cytometry and Cell Sorting Laboratory are shared resources partially supported by Laura and Isaac Perlmutter Cancer Center Support Grant P30CA016087 from National Institutes of Health The National Cancer Institute (NIH-NCI). We would like to thank Stephen T. Yeung and Ze Chen for assistance with manuscript preparation.

