## Supplemental Figures 1-4 for "A role for BCL6 in maintaining CX3CR1^+^ CD4^+^ T cells during helminth infection"

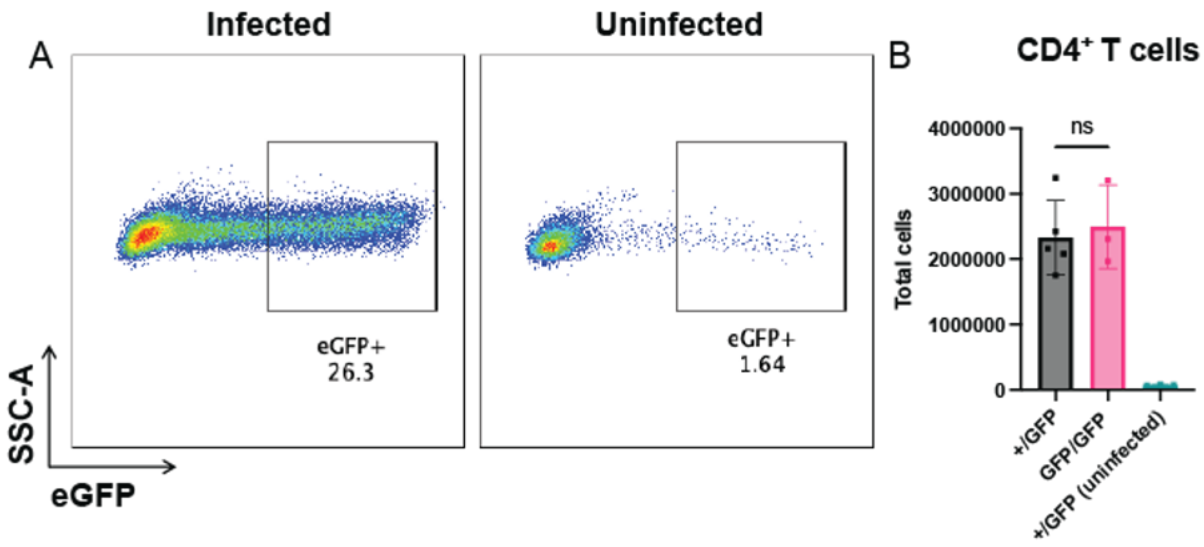


**Supplementary Figure 1. CX3CR1 is dispensable for CD4$^+$ T cell accumulation in the liver.** **(A)** Representative flow cytometry plot of eGFP expression in CD4$^+$ T cells isolated from the liver of CX3CR1$^{+/eGFP}$ mice infected with *S. mansoni*. **(B)** Quantification of eGFP$^+$ CD4$^+$ T cells in livers of mice infected with *S. mansoni*. Cells were gated on singlet cells, live cells, CD45$^+$ cells, CD11b$^{NEG}$ cells, CD3$^+$ cells, CD8$^{NEG}$, CD4$^+$ cells. N = 3-5 mice/group. Error bars represent SEM. Students *t* tests were performed to determine significance. *P<0.05; **P<0.01; ***P<0.001; ****P<0.0001.

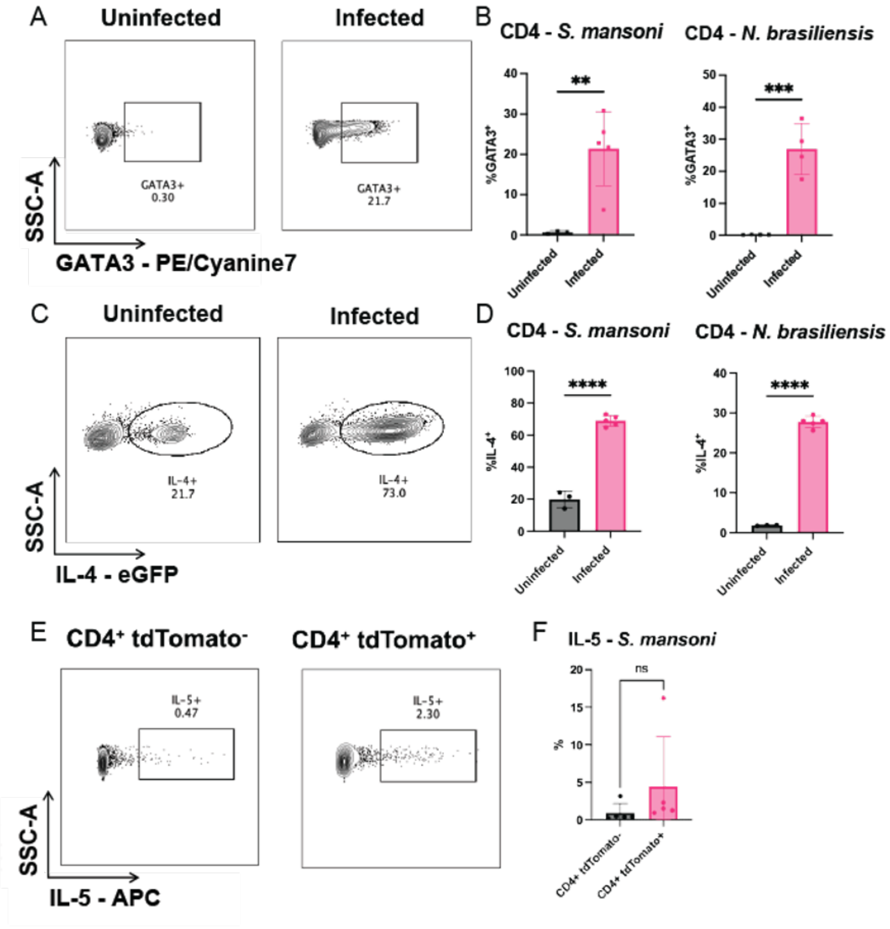


**Supplementary Figure 2. Percentage of total CD4$^+$ T cells with T$_H$2 phenotype increases due to infection. (A)** Representative flow cytometry plots of GATA3 expression on total CD4$^+$ T cells isolated from the liver of mice infected with *S. mansoni*. **(B)** Percentage of total CD4$^+$ T cells in liver of *S. mansoni* infected mice and lungs of *N. brasiliensis* infected mice that express GATA3 **(C)** Representative flow cytometry plots of IL-4 expression in total CD4$^+$ T cells isolated from the liver of 4get mice infected with *S. mansoni*. **(D)** Percentage of total CD4$^+$ T cells in liver of *S. mansoni* infected 4get mice and lungs of *N. brasiliensis* infected 4get mice that express IL-4. **(E)** Representative flow cytometry plots of IL-5 expression in tdTomato$^+$ and tdTomato$^-$ CD4$^+$ T cells in mice infected with *S. mansoni* and **(F)** quantification of percentage. Cells were gated on singlet cells, live cells, CD45$^+$ cells, CD11b$^{NEG}$ cells, CD3$^+$ cells, CD8$^{NEG}$, CD4$^+$ cells. N = 4-5 mice/group. Error bars represent SEM. Students *t* tests were performed to determine significance. *P<0.05; **P<0.01; ***P<0.001; ****P<0.0001.

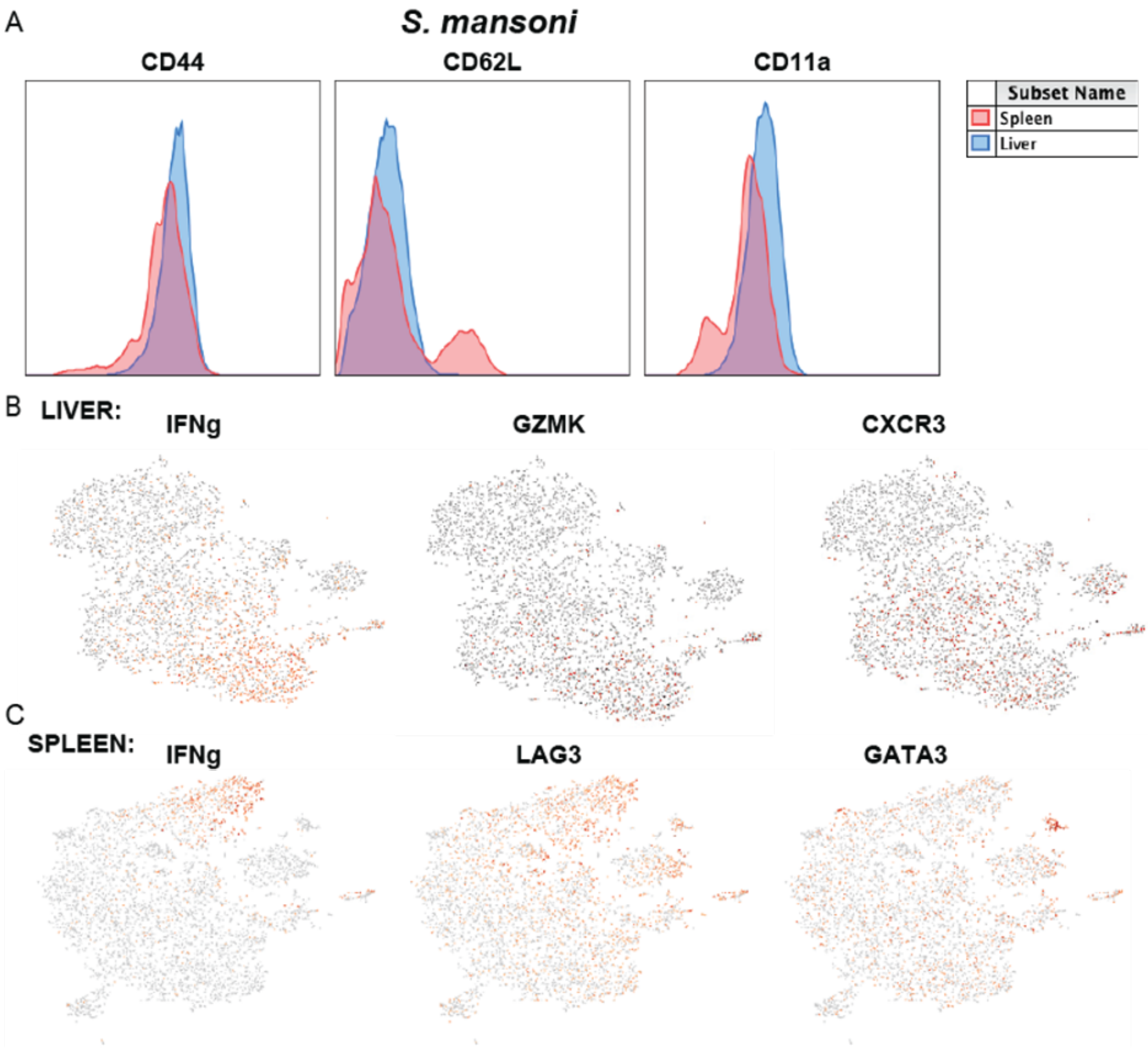


**Supplementary Figure 3. CD4$^+$ tdTomato$^+$ T cells in the spleen display a distinct phenotype.** **(A)** Overlayed histograms of activation marker expression in both liver and spleen from mice infected with *S. mansoni* at 8 weeks post-infection. **(B)** UMAP clustering of tdTomato$^+$ CD4$^+$ T cells from the liver of mice infected with *S. mansoni* with selected genes highlighted. **(C)** UMAP clustering of tdTomato$^+$ CD4$^+$ T cells from the spleen of mice infected with *S. mansoni* with selected genes highlighted. For analysis, cells were gated on singlet cells, live cells, CD45$^+$ cells, CD11b$^{NEG}$ cells, CD3$^+$ cells, CD8$^{NEG}$, CD4$^+$ cells. For sorting/scRNA-seq, cells were sorted as described in methods.

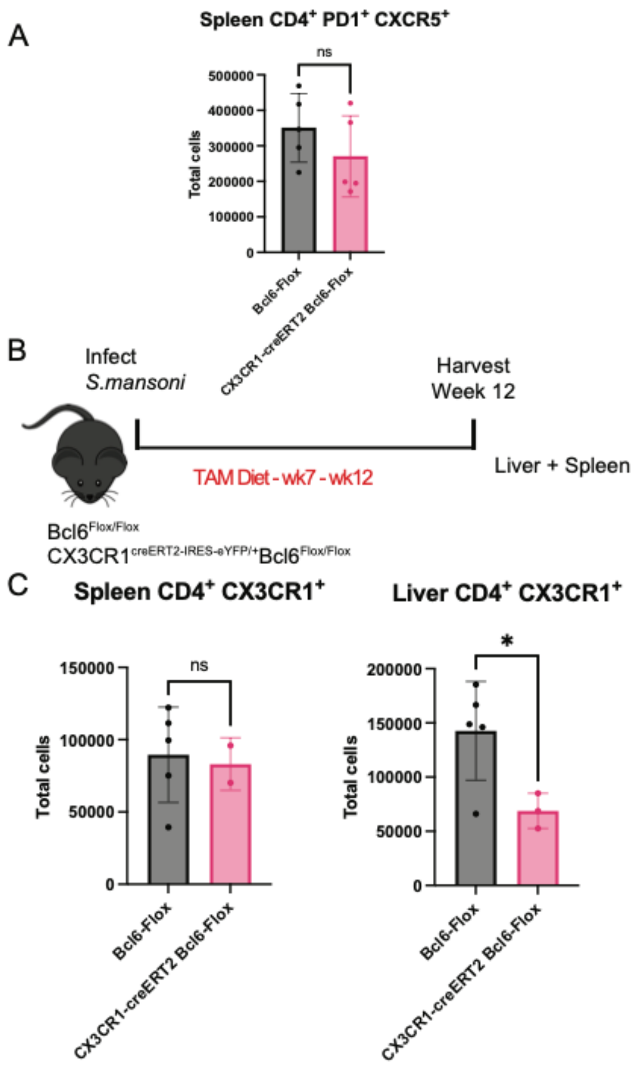


**Supplementary Figure 4. CX3CR1-specific deletion of BCL6 reduces accumulation of CD4$^+$ CX3CR1$^+$ cells in the liver during *S. mansoni* infection (A)** Quantification of CD4$^+$ PD1$^+$ CXCR5$^+$ T cells in the spleen of Bcl6$^{Flox}$ and CX3CR1$^{CreERT2}$ Bcl6$^{Flox}$ mice infected with *N. brasiliensis*. Cells were gated on singlet cells, live cells, CD45$^+$ cells, CD11b$^{NEG}$ cells, CD3$^+$ cells, CD8$^{NEG}$, CD4$^+$, PD1$^+$ CXCR5$^+$ cells. **(B)** Experimental design of experiment to conditionally delete Bcl6 in CX3CR1$^+$ cells **(C)** Quantification of CX3CR1$^+$ CD4$^+$ T cells in the spleen and liver of Bcl6$^{Flox}$ and CX3CR1$^{CreERT2}$ Bcl6$^{Flox}$ mice infected with *S. mansoni*. Cells were gated on singlet cells, live cells, CD45$^+$ cells, CD11b$^{NEG}$ cells, CD3$^+$ cells, CD8$^{NEG}$, CD4$^+$ cells. N = 2-5 mice/group. Error bars represent SEM. Students *t* tests were performed to determine significance. *$P<0.05$; **$P<0.01$; ***$P<0.001$; ****$P<0.0001$.
